# Recombinant Fc-fusion vaccine of RBD induced protection against SARS-CoV-2 in non-human primate and mice

**DOI:** 10.1101/2020.11.29.402339

**Authors:** Shihui Sun, Lei He, Zhongpeng Zhao, Hongjing Gu, Xin Fang, Tiecheng Wang, Xiaolan Yang, Shaolong Chen, Yongqiang Deng, Jiangfan Li, Jian Zhao, Liang Li, Xinwang Li, Peng He, Ge Li, Hao Li, Yuee Zhao, Chunrun Gao, Xiaolin Lang, Xin Wang, Guoqiang Fei, Yan Li, Shusheng Geng, Yuwei Gao, Wenjin Wei, Zhongyu Hu, Gencheng Han, Yansong Sun

**Author notes:** Corresponding to: Yuwei Gao, WenjinWei, Zhongyu Hu, Gencheng Han, Yansong Sun. These authors contributed equally (Shihui Sun, Lei He, Zhongpeng Zhao, Hongjing Gu, Xin Fang, Tiecheng Wang, Xiaolan Yang, Shaolong Chen, Yongqiang Deng).

## Abstract

The severe acute respiratory syndrome coronavirus-2 (SARS-CoV-2) continues to infect people globally. The increased COVID-19 cases and no licensed vaccines highlight the need to develop safe and effective vaccines against SARS-CoV-2 infection. Multiple vaccines candidates are under pre-clinical or clinical trails with different strengths and weaknesses. Here we developed a pilot scale production of a recombinant subunit vaccine (RBD-Fc Vacc) with the Receptor Binding Domain of SARS-CoV-2 S protein fused with the Fc domain of human IgG1. RBD-Fc Vacc induced SARS-CoV-2 specific neutralizing antibodies in non-human primates and human ACE2 transgenic mice. The antibodies induced in macaca fascicularis neutralized three divergent SARS-CoV2 strains, suggesting a broader neutralizing ability. Three times immunizations protected Macaca fascicularis (20ug or 40ug per dose) and mice (10ug or 20ug per dose) from SARS-CoV-2 infection respectively. These data support clinical development of SARS-CoV-2 vaccines for humans. RBD-Fc Vacc is currently being assessed in randomized controlled phase 1/II human clinical trails.

**Summary:** This study confirms protective efficacy of a SARS-CoV-2 RBD-Fc subunit vaccine.

## Introduction

The COVID-19 pandemic, caused by a novel Coronavirus SARS-CoV-2, has infected more than 60 million individuals and killed more than 1.4 million people globally^1^. Because of the high prevalence, longer incubation, no preexisting immunity to SARS-CoV-2 in human and even no effective treatment or drugs currently available, it is urgent to develop safe and efficient vaccines to control and prevent the further dissemination of SARS-CoV-2.

SARS-CoV-2 showed similar but distinct genome composition of SARS-CoV and MERS-CoV, another two coronaviruses in the genus Beta coronavirus with zoonotic origin. So, by the knowledge from SARS and MERS vaccines development path, several vaccine candidates, such as DNA vaccines, RNA vaccines, viral vector vaccines, recombinant subunit protein vaccines, inactivated virus vaccines are all developed^2,3^ and at least three inactivated SARS-CoV-2 vaccines are in phase III clinical trials^4^. Compared with other vaccines, recombinant subunit protein vaccine shows many advantages such as safety, higher cost efficiency and easy to be handled^3^. The main concern about the recombinant protein vaccine is that whether it can induce enough neutralizing antibodies and provide protection. Fortunately, recombinant subunit vaccine targeting SARS-CoV-2 RBD induced higher neutralizing antibodies without evident antibody dependent enhancement effects (ADE) ^5^ and may protect animals from SARS-CoV-2 attack ^6-7^.

SARS-CoV-2, homologous to SARS-CoV and the Middle East respiratory syndrome coronavirus, also contains four structural proteins, including spike (S), envelope (E), membrane (M), and nucleocapsid (N) proteins. Among them, S protein plays the most important role in viral attachment, fusion and entry, and induces neutralizing antibodies with virus infection, so serves as a target for development of vaccines. The immunogenicity of SRAS-CoV-2 RBD, lies in S1 subunit by binding to the receptor ACE2 and is crucial in mediating viral entry into host cells, has been studied to induce neutralizing antibodies, which has been verified in samples from patients recovered from COVID-19 ^8^. These data indicate that RBD-based subunit proteins vaccine is a competitive candidate.

The Fc fusion protein is recently used as an important backbone for drug development, owing to its advantages including an expedient for rapid purification, longer half-life, increasing immunogenicity of target antigens and eliciting neutralizing antibody response^9^. In addition, Fc promote the correct folding of fusion protein and to enhance binding to antigen-presenting cells (APCs) and cell lines expressing Fc receptors (FcR)^10,11^. For example, Enbrel, TNF Receptor-Fc fusion protein used for the treatment of rheumatoid arthritis or other human diseases, is a therapeutic biological drug without any known side-effect ^12^. We have previously developed Fc-fused protein vaccines against MERS, SARS-CoV and H5N1 influenza, and found that Fc fused protein is more immunogenic than those lacking fused Fc^13-15^.

In this study, we designed a similar recombinant subunit vaccine (RBD-Fc Vacc) with the Receptor Binding Domain of SARS-CoV-2 S protein fused the Fc domain of human IgG1. The RBD-Fc Vacc showed immunogenicity and protective efficacy in non-human primate and human ACE2 transgenic (hACE2 Tg) mice with broader neutralizing ability against divergent SARS-CoV-2 virus strains. This novel COVID-19 subunit vaccine is currently being evaluated in phase I/II clinical trials. And for our best knowledge, this is the first Fc fusion protein-based vaccine tested in clinical trials.

## Results

### Construction and antigenicity of RBD-Fc Vacc

Here a dimer-based RBD-Fc Vacc against SARS-CoV-2 was developed by fusing RBD (aa331-524) with human IgG1 Fc. The predicted dimer structure is shown in Fig 1a (left panel) by protein structure prediction server version 3.0 ^16^: Two RBD domains are fused through Fc fragment to form the Y-shaped structure. The RBD-Fc fusion protein was expressed in mammalian CHO cells and the antigenicity and the dimer conformation were identified by western blot analysis using anti-serum from COVID-19 recovered patients and commercial antibody respectively. (Fig.1a right panel). The expressed protein is about 120 kDa under non-reduced condition and 60 kDa after reduced. The calculated molecular weight of RBD-Fc protein is about 95.9 kDa while the purified recombinant protein appears to be 120kDa on the western blot owning to protein glycosylation. Three N-glycosylation sites on asparagine, one O-glycosylation on serine and one O-glycosylation on threonine were identified using mass spectrum (Fig.1b up panel). These identified glycosylation sites were further mapped on the complex structure of SARS-CoV-2 RBD-Fc bound to ACE2 predicted by ZDOCK server^17^. The glycosylation sites were located on the RBD core subdomain and were found to be distant from the area bound to ACE2 (Fig. 1b down panel), indicating that the decorated glycopeptides may not interfere with receptor recognition.

**Fig.1.**
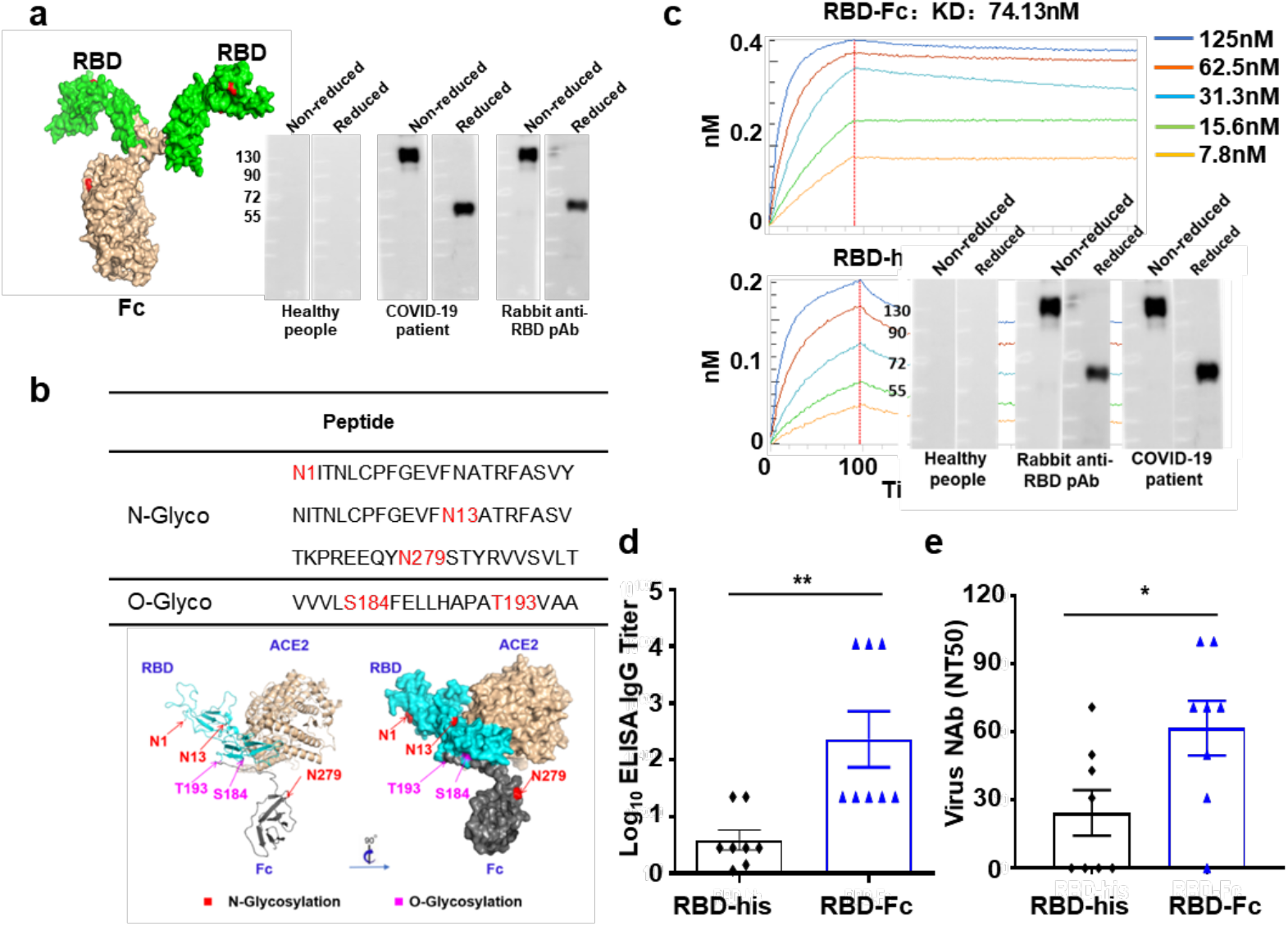
SARS-Cov2 RBD-Fc vaccine Design. **a)** Two RBD domains are fused through Fc fragment to form the Y-shaped structure via protein structure prediction server version 3.0 (left panel). RBD-Fc protein expressed from CHO cells were identified by western blot under reduced and non-reduced condition using both anti-serum from COVID-19 recovered patients and commercial antibody (right panel). b) N-glycosylation and O-glycosylation sites identified using mass spectrum (up panel). The docking between ACE-2 and RBD-Fc predicted by ZDOCK server. An overview of the glycosylation sites illustrated based on the solved complex structure of SARS-CoV-2 RBD-Fc bound to ACE2 (PDB code: 1R42). The identified sites, colored red for N-glycosylation, purple for O-glycosylation are shown as spheres and labeled. The right panel (surface representation) was generated by rotating the structure in the Left panel (cartoon representation) around a vertical axis for about 90° (lower panel). c) The real-time binding profile between our purified RBD-Fc protein and ACE2 characterized by SPR Biacore. d&e) Balb/C mice were immunized with RBD-his or RBD-Fc (10ug/mouse) in the presence of aluminum at d0 and d14. Two weeks post last vaccination the serum were collected ELISA assay shows the SARS-CoV-2 RBD-specific IgG titers (d). SARS-CoV-2 neutralization assay shows the NT50 (e).

The binding of RBD-Fc protein with ACE2 was then confirmed via surface plasmon resonance (SPR) Biacore and potent interactions were observed in concordance with a previous study^7^. Results from the SPR assay demonstrated that RBD-Fc protein bound to receptor hACE2 with much higher avidity compared with RBD-His monomer (74.13nM vs. 8.26nM), indicating the better conformation of our RBD-Fc protein (Fig.1c). The immunogenicities of RBD-Fc or RBD-His protein were tested in BALB/c mice after two immunizations using aluminum as the adjuvant. Compare to those from RBD-His vaccinated mice, serum from RBD-Fc vaccinated mice showed significantly higher RBD specific IgG antibody titers (with median peak of 1/40765 vs 1/7845) and live virus neutralizing antibody titers (with median peak of 1/61.8 vs. 1/24.4) (Fig.1d&e). These data suggested that RBD-Fc fusion protein is more antigenic than monomer RBD protein.

### The immunogenicity of RBD-Fc Vacc in Macaca fascicularis and hACE2-Tg mice

We evaluated the immunogenicity of RBD-Fc Vacc in 3-year-old Macaca fascicularis, a non-human primate model that was susceptible to SARS-CoV-2 infection^18-20^. Two groups of Macaca fascicularis (n=5/each group) were immunized with 20ug or 40ug of RBD-Fc Vacc via intramuscular administration at 0 days(prime),14 days (1^st^ boost) and 28 days (2^nd^ boost) and the equal number of macaques (n=5) received PBS injection as the sham control. No inflammation or other adverse effects were observed during the observation period. Serum were collected at each time point as shown in Fig.2a to examine RBD specific antibodies as well as neutralizing antibodies using ELISA or neutralization assay. High level of RBD-specific immunoglobulin G (IgG) were induced quickly in the serum of RBD-Fc Vacc immunized Macaca fascicularis up to a median peak of 1/17399 in 20ug group and of 1/41900 in 40ug group at day 28 (two weeks post 1^st^ boost) (Fig.2b). Although there is a moderate increase for RBD-specific IgG titers in serum collected at day 42 compare to those at day 28 (Fig.2b), there is a much higher NT50 in sera from three dose-RBD-Fc Vacc-immunized macaca fascicularis (day 42, boost twice) compare to those from two dose-RBD-Fc Vacc-immunized animals (d28, boost once) (Fig.2c), which indicated that three dose of immunization may induce a much better protective response. The variability in immunogenicity in this study facilitated an analysis of correlations of different examining methods. NAb titers measured by live virus neutralization assay correlated with RBD specific IgG titers (Fig.2d). Because of the novel epidemic SARS-CoV-2 strains with specific mutations was recorded ^20-21^, we further tested whether Macaca fascicularis sera post RBD-Fc vaccination could cross neutralize different epidemic strains of SARS-CoV-2. Here three epidemic strains BJ08, BJ05 and BJ01, which shared the same RBD sequence as our RBD-Fc vaccine^22^, were used to test the cross neutralization effects. All the sera from RBD-Fc Vacc immunized Macaca fascicularis showed similar neutralizing capability against three SARS-CoV-2 epidemic strains, and there was no significant difference in NT50 titers (Fig.2e). In addition, the NT50 in sera from non-human primates receiving RBD-Fc Vacc immunization was comparable to those from 23 COVID-19 convalescent patients’ serum. (Fig.S1). Our results demonstrated that three immunization of RBD-Fc Vacc have induced high levels of antibodies with broad neutralizing capabilities against SARS-CoV-2 in non-human primates.

**Fig.2.**
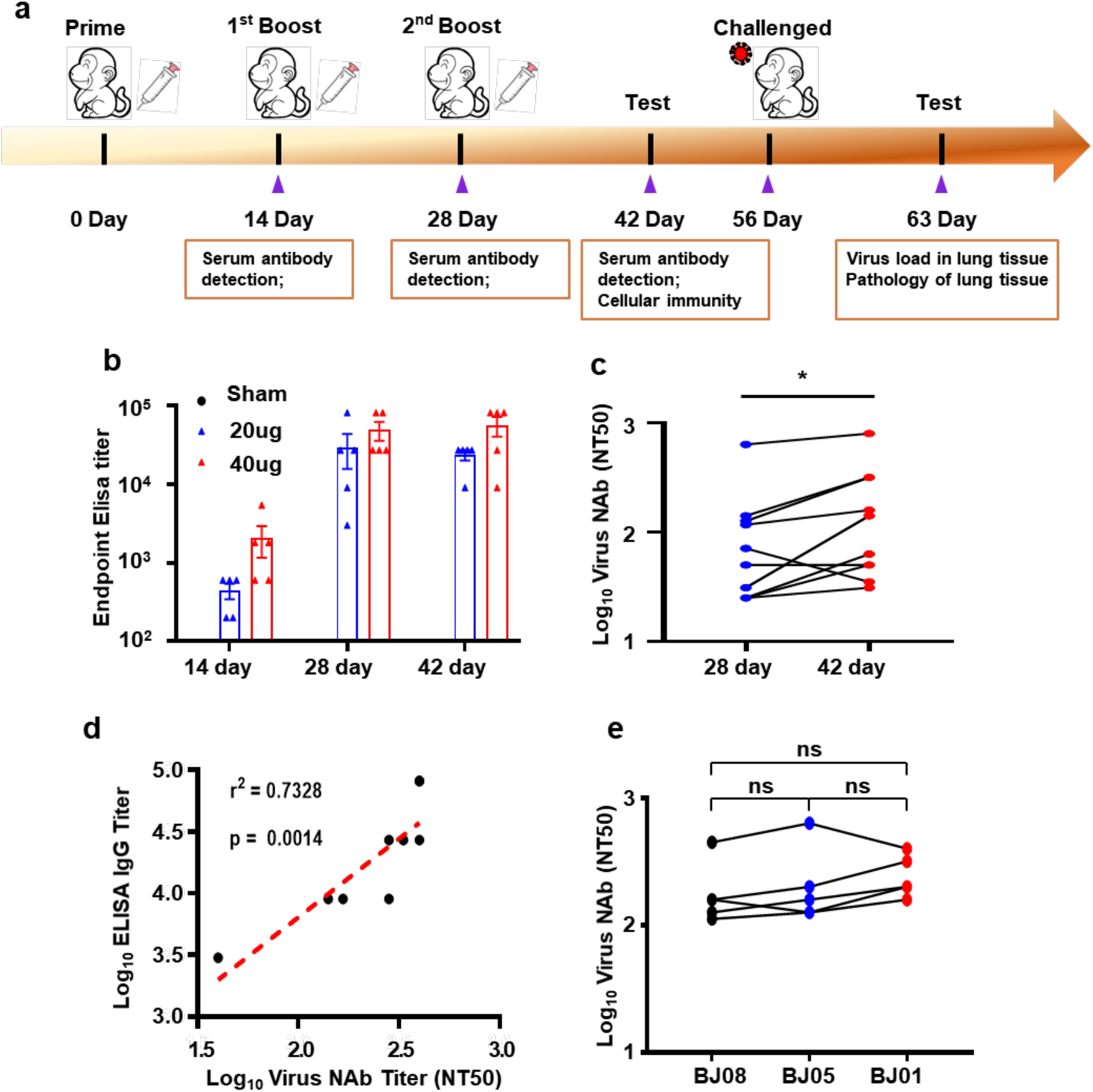
SARS-Cov2 RBD-Fc Vacc elicit strong humoral immunity in macaca fascicularis. a) Schematic diagram of immunization, sample collection and challenge schedule. b). Macaca fascicularis (n=5) were immunized on day 0, day 14, d28 with 20ug and 40ug doses of RBD-Fc Vacc or PBS and the serum were collected at the indicated time. The SARS-CoV-2 RBD specific IgG were examined by ELISA. c) Neutralizing antibodies were determined by microneutralization assay using the SARS-CoV-2 (NT50). d). Correlation analysis of antibody titers tested by ELISA and microneutralization assay. P and R values reflect two-sided Spearman rank-correlation tests. e) Serum cross neutralization against SARS-CoV-2 epidemic strains in RBD-Fc immunized macaca fascicularis. NT50 against the SARS-CoV-2 epidemic strains (BJ08, BJ05, BJ01) were performed using macaca fascicularis sera collected at 21 days post third immunization. Data are analyzed by one-way ANOVA with multiple comparison tests. (n.s., not significant).

The immunogenicity of RBD-Fc Vacc was also confirmed in hACE2 Tg (n=8). The immunizing schedule in mice was in accordance with Macaca fascicularis. RBD-specific immunoglobulin G (IgG) and the neutralizing antibodies developed quickly in the serum of vaccinated hACE2-Tg mice. And no significant difference was found between 10ug and 20ug RBD-Fc Vacc induced humoral immune response (Fig. S2).

To compare the cellular immune response, IFN-γ ELISPOT assay and flow cytometry (d42) were performed in Macaca fascicularis at 2 weeks post 2^nd^ boost. There are comparable S1 or RBD specific IFN-γ response among splenocytes from non-vaccinated or from RBD-Fc Vacc vaccinated macaca fascicularis (Fig. S3a), and the percentages of S1-specific IL-4^+^CD4^+^ T cells, IL-4^+^CD8^+^ T cells; TNF-a^+^CD4^+^T cells, TNF-a^+^CD8^+^ T cells were at the same level (Fig. S3b), suggesting that RBD-Fc Vacc mainly induces humoral immune response in vaccinated animals.

### Protective efficiency against SARS-CoV-2 challenge in Macaca fascicularis and in hACE2-Tg mice

We next evaluated whether RBD-Fc Vacc induces protective immunity against SARS-CoV-2 in Macaca fascicularis. Four weeks after the third injection of 40ug RBD-Fc Vacc (d56), macaques were challenged with 10^6^ TCID50/mL SARS-CoV-2 as follows: 2 mL inoculation by the intratracheal route, 1 mL by the intranasal route and 0.2 mL by the intraocular route. The substantial fraction of viral RNA which represent input challenge virus in Nasal, throat and anal swabs were examined at different times following challenge by quantitative real-time reverse transcription-PCR (qRT-PCR). Peak viral loads occurred variably following challenge. All control Macaca fascicularis showed excessive copies of viral genomic RNA in the nasal, throat, anal and lung by day 2-6 post inoculation (Fig.3a, b and Table S1) and severer interstitial pneumonia (Fig.3c). By contrast, all vaccinated Macaca fascicularis were largely protected against SARS-CoV-2 infection with much lower or absence of viral RNA copies and very mild histopathological changes in a few lobes of lung. These data demonstrated that RBD-Fc Vacc induced potent immune response and efficiency protected Macaca fascicularis from SARS-CoV-2 attack.

**Fig.3.**
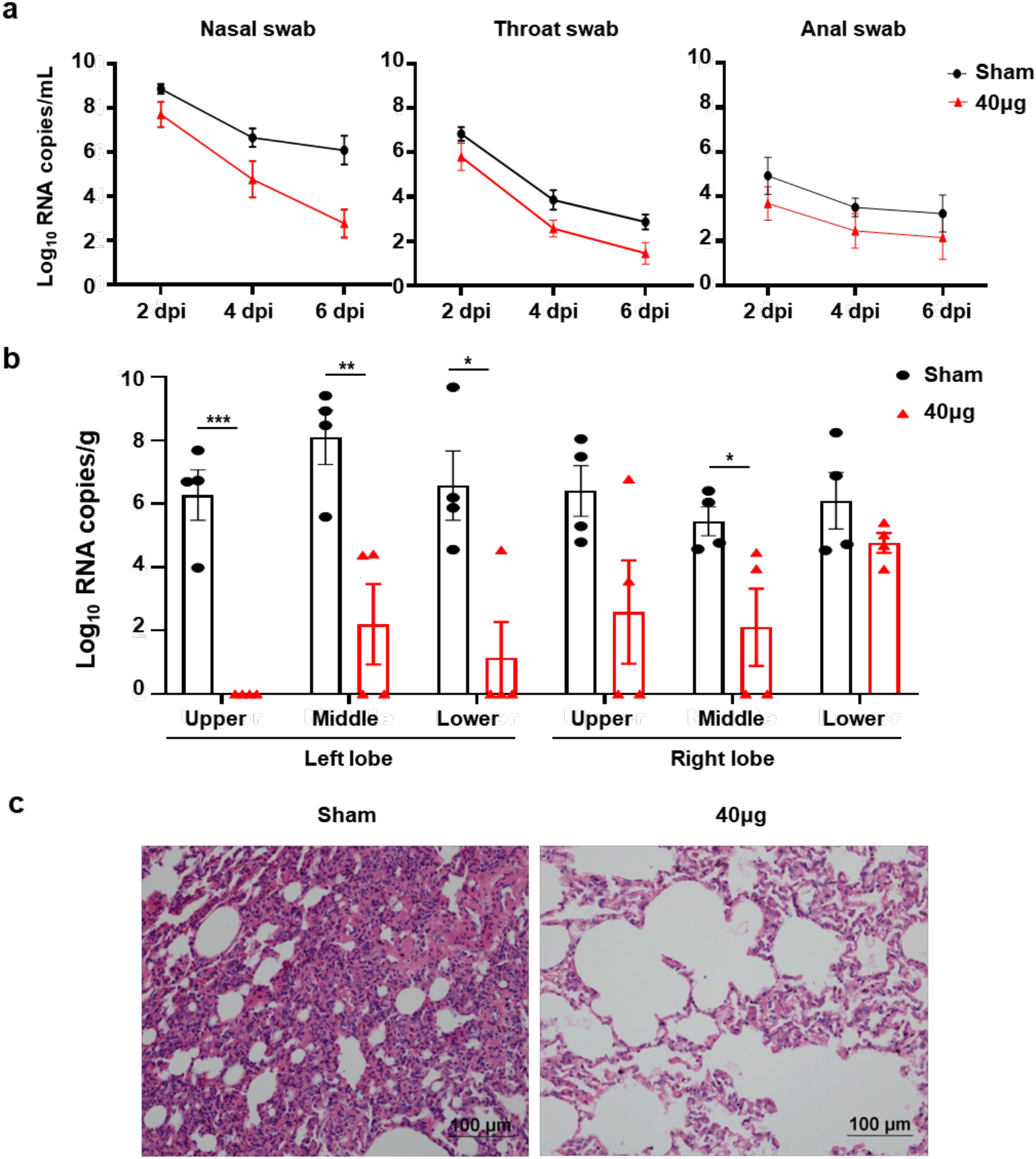
Protective efficacy of RBD-Fc Vacc in macaca fascicularis. **a)**. Viral load of nasal, throat, and anal specimens collected from the inoculated immunized macaca fascicularis (n=4) at day 2, day 4 and day6 pi were monitored. b) Viral loads in various lobes of lung tissue from the inoculated macaca fascicularis (n=4) at day 7 post infection were measured. All data presented as mean +SEM from two independent experiments. c) Histopathological examinations in lungs from the inoculated macaca fascicularis (n=4) at day 7 post injection. Lung tissue was collected and stained with hematoxylin and eosin.

The protective efficiency of RBD-Fc Vacc against SARS-CoV-2 was also confirmed in hACE2-Tg mice. Four weeks after the second boost immunization, mice were challenged intranasally with 1.8 x 10^7^ PFU/ml SARS-CoV-2. Five days following challenge, viral RNA levels in lung were assessed by RT-PCR. RBD-Fc Vacc immunization in hACE2-Tg mice lead to a median reduction of 2.10 Log_10_RNA copies/g and 2.13 in 10ug and 20ug group respectively (Fig.S4a). In addition, all vaccinated animals were largely protected against SARS-CoV-2 infection with very mild histopathological changes in the vaccinated animals, while moderate interstitial pneumonia in sham group. (Fig.S4b).

### Immune correlates of vaccine-induced protection

The variability in protective efficiency in this study facilitated an analysis of immune correlations of protection. The live virus neutralization antibody titers at day49 inversely correlated with viral RNA in lung of hACE2-Tg mice (Fig. 4a). In addition, the pseudovirus neutralization antibody titers and the IgG titers at day 49 also indicated inversely correlation with viral RNA in lung of hACE2-Tg mice (Fig. 4b&c). The vaccine induced immune response and protection also showed some inverse correlation in Macaca fascicularis, however, due to the limitation of sample numbers, the data cannot be statistically analyzed. These data demonstrated that serum IgG titers and neutralization antibody titers elicited by RBD-Fc may be immune correlates of protection against SARS-CoV-2 infection. The data also suggested that these parameters would be simple and useful benchmark for clinical development of SRAS-CoV-2 vaccine.

**Fig.4.**
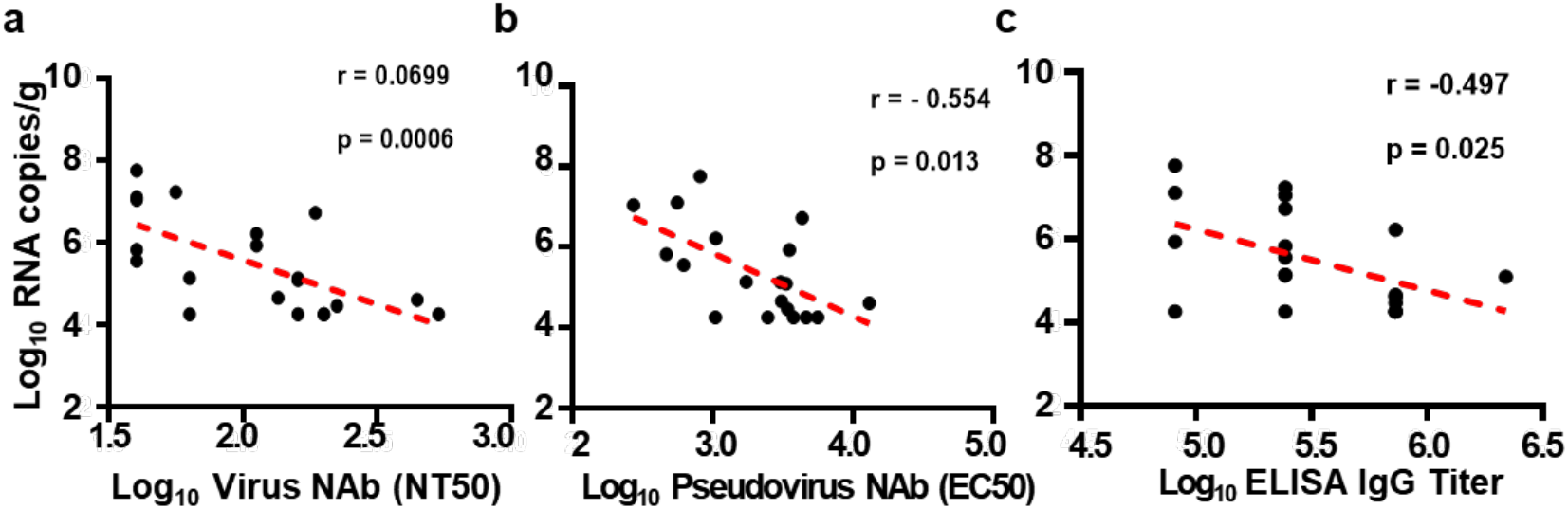
Correlations of SARS-CoV-2 NAb titers (a), pseudovirus titers (b), and RBD protein specific IgG titers (c) prior to challenge with log peak mRNA copies/g in lungs following challenge in hACE2-Tg mice. Red lines reflect the best-fit relationship between these variables. P and R values reflect two-sided Spearman rank-correlation tests.

## Discussion

The serious pandemic of current COVID-19 and the increasing numbers of death worldwide necessitate the urgent development of a SARS-CoV-2 vaccine. The safety and efficiency are essential for vaccine development at both stages of preclinical studies and clinical trials. In this study, we generated a new recombinant vaccine by fusioning SARS-CoV-2 RBD with the Fc domain of human IgG1 and assessed the protective efficacy against SARS CoV-2 challenge in nonhuman primates (macaca fascicularis) and hACE2-Tg mice respectively. Compared to the excessive copies of SARS-CoV-2 RNA and severe lung damage in non-immunized sham groups, all vaccinated macaca fascicularis were largely protected against SARS-CoV-2 infection with absence or much lower of viral RNA copies and non-or very mild histopathological changes in a few lobes of lung. It is also encouraging that our candidate RBD-Fc vaccine candidate induce effective neutralizing activity in hACE2-Tg mice. Most importantly, serum from vaccinated macaca fascicularis can neutralize different SARS-CoV-2 strains, demonstrating the wide spectrum of neutralizing activity induced by RBD-Fc Vacc.

By the knowledge and vaccine platform from SARS and MERS, several vaccines have been developed rapidly within a very short time. Inactivated vaccine is a traditional vaccine platform with rapid development and conceptual safety, but demands more strict and precise process in manufacturing which may determine a disease outbreak^3^. The new-generation vaccines, including vector-based vaccines and recombinant protein vaccines, are a relatively safety profile because of the selection of a specific antigen or antigens instead of the whole pathogen^23^. In addition to the concerns that viral genome may integrate into host genome^3^, scaling-up of viral production capacity is also a bottleneck of viral vector manufacturing^24^. Therefore, the current challenges of virus vector-based vaccines still lie in a good balance between high recovery yield and impurity clearance and the reduction of cost^25^. Other vector based vaccine including DNA and mRNA vaccines are facing challenges such as integration of foreign DNA into the host genome may cause abnormalities in the cell^3^ and maintaining of vaccines in a harsh cold chain^26-27^. Recombinant protein vaccine expressed in CHO system can maintain the right glycosylation and is an economic way to make enough vaccines for a large-scale population. Especially recombinant subunit protein vaccine is the most safety way as it does not have any live component of the viral particle^3^.

The most effective way to identify and deploy efficacious SARS-CoV-2 vaccines with acceptable safety profiles is through carefully designed. SARS-CoV-2 virus is homologous to SARS-CoV and the Middle East Respiratory Syndrome Coronavirus, which have been studied for years. Similar to SARS-CoV and MERS, S protein of SARS-CoV-2 has been identified as a key component triggering protective immunity. Previous studies in SARS-CoV vaccine development, liver damage was observed in the animal model when the full length S protein was used as the vaccine antigen^28^. Therefore, using S protein fragment, such as the receptor-binding domain (RBD), as vaccine antigen might be a safer choice for COVID-19 vaccine candidates. Sequencing data have showed that RBD is highly conserved among ten pandemic SARS-CoV-2 strains worldwide. Our data showed that serum from RBD-Fc Vacc immunized Macaca fascicularis neutralized three representative SARS-CoC-2 strains in microneutralization assay (Fig.3d). RBD-Fc-Vacc-elicited serum IgG titers and NAb titers also correlates negatively with the virus copies of immunized hACE2-Tg mice, demonstrating IgG and NAb titers as an immune correlate of protection. These data suggested that RBD is an excellent target for SARS-CoV-2 vaccine.

The antibody mediated enhancement effects (ADE) is the most important safety concerns in the development of vaccines. A recent report also showed that nonhuman primates infected with SARS-CoV-2 are protected from reinfection with the virus^29^, which indicated that virus infection induced a protective immune response. Reports showed that non-neutralizing antibodies is likely to promote more efficient ADE than neutralizing antibodies^30^. It has been noted that non-neutralizing coronavirus antibodies may cause ADE in feline infectious peritonitis. Such efforts have prompted investigators to focus on the RBD as a lead vaccine candidate while remove potential ADE-promoting S protein epitopes outside the RBD^13-31^. Indeed, a recent data showed that RBD of SARS-CoV-2 raised potently neutralizing antisera in immunized rats and these sera without ADE^5^. Our reports showed that RBD-Fc Vacc immunized macaca fascicularis and mice were protected from SARS-CoV-2 attack without evident side effects, which suggested that RBD is a safe and efficient target for vaccine use.

Fusion proteins based on immunoglobulin Fc domain have received considerable attention over the past two decades because of its capability to promote protein expression, purification, and improve the immunogenicity of the fused proteins. Many therapeutic biological drugs based on Fc protein fragment had been used for treatment of human diseases^32^. The application of Fc-fusion protein as vaccine delivery platform was first reported in 1989 in which gp120-Fc was used as a potential candidate for AIDS therapy^33^. Although there is no FDA approved Fc-fusion vaccine in clinic, the explosion of vaccine development fused with Fc is active and ongoing. A number of studies have been initiated on the development of vaccines against Ebola^34^, HIV^35^, influenza^36^, SARS-CoV^13^, tuberculosis^37^ as well as RSV^38^, HNV^39^. Currently, although other options, such as IgG3, IgA, and IgM are being explored, all commercial therapeutics use the Fc domain from human IgG1^40^. In addition, the ability of Fc to improve immune response has been confirmed in the studies of vaccine designation fused with RBD of H5N1, SARS-CoV and MERS-CoV^13,14,41^. Here, we found that Fc fragment enhances the immunogenicity of RBD as RBD-Fc induced much higher titer of S1 protein specific IgG antibodies and neutralization antibodies than RBD-His (Fig.1d&e), which further demonstrated that Fc-fusion protein is a promising vaccine delivery platform.

In summary, our data demonstrated that RBD-Fc Vacc, with adjuvant of aluminum, induced a robust humoral immune response and protected non-human primate and hACE2 Tg mice from SARS-CoV-2 infection. Importantly, our data demonstrate the immune correlates of protection and protective efficacy and support the immunogenicity of the RBD-Fc Vacc candidate in clinical use. For the best of our knowledge, this is the first Fc fusion protein-based vaccine tested in clinical trials. Further research will need to address the important questions of the durability of protective immunity and the optimal vaccine strategy for a SARS-CoV-2 vaccine for humans.

## Materials and Methods

### Ethics statement

All animal studies were performed in strict accordance with the guidelines set by the Chinese Regulations of Laboratory Animals and Laboratory Animal-Requirements of Environment and Housing Facilities. All animal procedures were reviewed and approved by the Animal Experiment Committee of Laboratory Animal Center, Academy of Military Medical Sciences (AMMS), China (Assurance Number: IACUC-DWZX-2020-040). Convalescent sera were collected from COVID-19 patients from Beijing YouAn Hospital, Capital Medical University with written informed consent.

### Gene cloning, protein expression and identify

The coding sequence for RBD-Fc region, which fused residues 331-524 from the spike protein of the SARS-CoV-2 and human IgG1-Fc together, were synthesized by Beijing JOINN Bilogic co.Ltd (Beijing, China) and then conducted into the GS expression vector ZY-CDMO. For protein expression, the plasmid was first transfected into CHO-K1 cells and then stable clones were isolated by the limiting dilution method in the presence of 50 μM of Methionine Sulfoximine (MSX). The cell culture supernatants were collected and analyzed by western blot analysis using a commercial antibody (Sino Biological Inc. Beijing, China) against SARS-CoV-2 RBD.

### Mass spectroscopy (MS) analysis to identify glycosylation sites

The recombinant RBD-Fc protein was purified using affinity chromatography and anion exchange chromatography. The purified RBD-Fc protein was then reduced by incubation with 10 mM Tris (2-carboxymethyl) phosphine for 20 min at 45°C in denaturation buffer (6 M guanidine hydrochloride, 20 mM Tris, 0.1 mM Na_2_-EDTA, pH 7.50). After cooling to room temperature, iodoacetamide was added to a final concentration of 60 mM and reaction was allowed to occur in the dark for 15 min. The protein was digested with Endoproteinase Lys-C (1:20 w/w) at 37°C overnight after buffer be replaced with the digestion buffer(20 mM Tris, pH 7.50). The digestion was stopped by formic acid solution and stored at 2~8°C until injection into the column.

The digested Peptide mixture was then separated on an Advance Bio Peptide Mapping column (2.1 ×150mm, 2.7μm) with flow rate of 0.2 mL/min and 75-min gradient (2-45% B) at 55°C. The mobile phase A was 0.1% formic acid in water, while mobile phase B contained 0.1% formic acid in acetonitrile. Liquid Chromatography (LC)-eluted peptides were detected by MS with an alternating low collision energy (4 V) and elevated collision energy (ramping from 25 to 40 V) ESI+ acquisition mode to obtain the precursor ions (MS) and their fragmentation data (MSE), respectively. A capillary voltage of 3.0 kV, source temperature of 120°C, cone voltage of 40 V, cone gas flow of 50 L/h were maintained during the analyses. The collected LC-MS data were processed by UNIFI software. Glycan structures were set as variable modifications. The mass tolerance for both precursors and fragments were set to less than 10 ppm.

### Surface plasmon resonance (SPR) analysis

SPR-based measurements were performed by Biacore T200 (GE Healthcare, Uppsala, Sweden), as described previously ^16^. Human ACE2-Avidin was captured to ~100RU on Sensor Chip Super Streptavidin. For kinetic analysis, RBD-Fc or RBD-his (Sino Biological Inc, Beijing, China) protein was run across the chip in a 2-fold dilution series (125, 62.5, 31.3, 15.6, 7.8nM), with another channel set as control. Each sample bound across the antigen surface was dissociated by HBS-EP (GE Healthcare) + running buffer for 300 s at a flow rate of 30 μL/min. Regeneration of the sensor chips was performed for 60 s using regeneration buffer (10mM Glycine-HCl pH 2.1). The association and dissociation rate constants ka and kd were monitored respectively and the avidity value KD was determined.

### Cells and Viruses

African green monkey kidney cell Vero (ATCC, CCL-81), were maintained in ulbecco’s minimal essential medium (DMEM; Thermo Fisher Scientific) supplemented with 10% fetal bovine serum (FBS; Thermo Fisher Scientific) and penicillin (100 /ml)-streptomycin (100 μg/ml) (Thermo Fisher Scientific).

Patient-derived SARS-CoV-2 isolates including 1) BetaCoV/Beijing/IMEBJ01/2020 (Patients originated from Beijing China; Accession No: GWHACAX01000000; Abbreviated as BJ01); 2) BetaCoV/Beijing/IMEBJ05/2020(Patients originated from Wuhan, China; Accession Nos: GWHACBB01000000. Abbreviated as BJ05);3) BetaCoV/Beijing/IMEBJ08/2020 (Patients originated from: Italy, Accession Nos: GWHAMKA01000000, Abbreviated as BJ08) were passaged in Vero cells and the virus stock was aliquoted and titrated to PFU/ml in Vero cells by plaque assay^22^. All experiments involving infectious SARS-CoV-2 were performed under Biosafety Level 3 facilities in AMMS.

### Animal immunization and challenge studies

Eight-week old human ACE2 transgenic mice (hACE2-Tg, obtained from National Institutes for Food and Drug Control, Beijing, China), Balb/C mice and 3-year old Macaca fascicularis (all from Laboratory animal center of the academic of the military medical sciences) were immunized with SARS-CoV-2 RBD-Fc Vacc as described in Fig.1a. Four weeks after the second boost immunization, macaca fascicularis were inoculated with 10^6^TCID50/ml BetaCoV(BJ01) as following: intratracheal 2ml, intranasal 0.5ml, intraocular 0.2ml; hACE2-Tg mice were inoculated intranasally with 40 μL of BetaCoV(BJ05). All mice or Macaca fascicularis were observed daily. On day 5 post infection (for mice), or on day 7 post infection (for Macaca fascicularis), eight mice in each group and four Macaca fascicularis in each group were sacrificed, and their lungs were removed for detection of viral load or were embedded for pathological analysis as described below.

### ELISA

ELISA was performed to detect SARS-CoV-2 RBD and S1-specific IgG antibodies in the immunized mouse or Macaca fascicularis sera. Briefly, ELISA plates were precoated with SARS-CoV-2 S1 or RBD protein (2.0μg/ml) overnight at 4°C and blocked with 3% BSA in PBS for 2 h at 37°C. Serially diluted sera were added to the plates and incubated for 45 min at 37°C. After four washes, the bound antibodies were detected by incubation with horseradish peroxidase (HRP)-conjugated anti-mouse IgG or anti-Macaca fascicularis IgG antibody (1:5,000, Thermo Fisher Scientific) for 30 min at 37°C. The reaction was visualized by addition of substrate 3,3’,5,5’-Tetramethylbenzidine (TMB) (Sigma, St. Louis, MO) and stopped by H2SO4 (1N). The absorbance at 450 nm was measured by a microtiter plate reader (Tecan, San Jose, CA).

### SARS-CoV-2 Pseudovirus based neutralization assay

The SARS-CoV-2 pseudovirus based neutralization assay was performed as described previously^42^. In brief, mice or Macaca fascicularis sera at 3-fold serial dilutions were incubated with 650 TCID50 of the pseudovirus for 1 hour at 37 °C, and then 20000 Huh7 cells were added into each well. DMEM was used as negative control. After 24h incubation in 37°C, the supernatant was then removed and luciferase substrate was added to each well followed by incubation for 2 minutes in darkness at room temperature. Luciferase activity was then measured using GloMax® 96 Microplate Luminometer (Promega). The 50% neutralization titer (NT50) was defined as the serum dilution at which the relative light units (RLUs) were reduced by 50% compared with the virus control wells.

### SARS-CoV-2 wild type virus neutralization assay

A micro-neutralization assay was carried out to detect neutralizing antibodies against SARS-CoV-2 infection. Briefly, mouse or Macaca fascicularis sera at 2-fold serial dilutions were incubated with SARS-CoV-2 (BetaCoV/BJ01); 100 TCID50) for 1 h at 37°C and added to Vero cells. The cells were observed daily for the presence or absence of virus-induced Cytopathic Effect (CPE) and recorded at 72 h. Neutralizing antibody titers were determined as the highest dilution of sera that can completely inhibit virus-induced CPE in at least 50% of the wells (NT50).

### Measurement of viral RNA

Tissue homogenates were clarified by centrifugation at 6,000 rpm for 6 min, and the supernatants were collected. Viral RNA (vRNA) was extracted using the QIAamp Viral RNA Mini Kit (Qiagen) according to the manufacturer’s protocol. vRNA quantification in each sample was performed by quantitative reverse transcription PCR (RT-qPCR) targeting the S gene of SARS-CoV-2. RT-qPCR was performed using One Step PrimeScript RT-PCR Kit (Takara, Japan) with the following primers and probes: CoV-F3 (5’-TCCTGGTGATTCTTCTTCAGGT-3’); CoV-R3 (5’-TCTGAGAGAGGGTCAAGTGC-3’); and CoV-P3 (5’-FAM-AGCTGCAGCACCAGCTGTCCA-BHQ1 −3’)

### IFN-γ ELISpot

PBMCs were collected from Macaca fascicularis and suspended in RPMI1640 media supplemented with 10% FBS and penicillin/streptomycin and processed into single cell suspensions. ELISpot assays were performed using the Macaca fascicularis IFN-γ ELISpot PLUS plates (MABTECH). 96-well ELISpot plates precoated with capture antibody were blocked with RPMI1640 medium overnight at 4 °C. 8 x 10^5^ PBMCs were plated into each well and stimulated for 48 h-72 h with S1 or RBD-his protein. The spots were developed based on manufacturer’s instructions. Medium and anti-CD3, anti-CD28 antibody were used for negative and positive controls, respectively. Spots were scanned and quantified by ImmunoSpot CTL reader. Spot-forming unit (SFU) per million cells was calculated by subtracting the negative control wells.

### Intracellular cytokine staining

Intracellular production of the cytokines including tumor necrosis factor alpha (TNF-a), and interleukin-4 (IL-4) was examined by flow cytometry. Briefly, PBMCs were harvested from Macaca fascicularis and stimulated with the SARS-CoV-2 S protein for 6 h at 37 °C, 5% CO_2_. Then cells were first stained with Percp conjugated anti-monkey CD4 (RM4-5, BD, 561115), Percp conjugated anti-monkey CD8a (RPA-T8, BD, 560662), at 4°C for 25 min. PE conjugated anti-monkey IL-4 (8D4-8, BD, 551774), and PE conjugated anti-monkey TNF-a (MAb11, BD, 557068) using the Cytofix/Cytoperm fixation/permeabilization solution kit (BD) according to the manufacturer’s instructions. Cell events were acquired using an FACS CANTO (BD), followed by FlowJo software (FlowJo LLC, Ashland, OR) analysis.

### Histopathological analysis

Five-or seven-days post virus challenge, animals were sacrificed and lung tissues were obtained, and paraffin-embedded in accordance with the standard procedure. Sections at 5 μm thickness were stained with hematoxylin and eosin (H&E), and examined by light microscopy. Lung tissue lesions were assessed according to the extent of denatured and collapsed bronchiole epithelial cells, degeneration of alveoli pneumocytes, infiltration of inflammatory cells, edema, hemorrhage, exudation and expansion of parenchymal wall. The semi-quantitative assessment was scored for the severity of lung damage according to the above criteria (0, normal; 1, mild; 2, moderate; 3, marked). The cumulative scores of the severity provided the total score per animal, and the average of six to eight animals from each group was taken as the total score for that group.

### Statistical analysis

Statistical analyses were carried out using Prism software (GraphPad). All data are presented as means ± standard error of the means (SEM). Statistical significance among different groups were calculated using the Student’s t test, Fisher’s Exact test, two-sided Spearman rank-correlation test, or Mann-Whitney test. *, **, and *** indicate P < 0.05, P < 0.01, and P < 0.001, respectively.

## Author contribution

S.S., L.H., Z.Z.,H.G., X.F.,W,T., X.Y., Y.D., S.C., J.L., T.W., J.Z.,L.L, X.L.,P.H.,G.L., H.L., C.G., X.L.,C.W., X.W., and G.F. performed experiments; S.S., L.H., H.G., Z.Z.,Y.L., S.G., W.W., Z.H., and G.H. analyzed data; Y.L., S.Y.(Yansong Sun) conceived the project. S.S., G.Y., H.G., Z.Z., G.H. Y.L., W.W., Z.H., and S.Y. (Yansong Sun) supervised the study and wrote the manuscript with the input of all co-authors.

## Acknowledgements

This article is in memory of Prof. Yusen Zhou for his contributions on the project conceive and article designation. We also thank Dr. X.D. Yu and Dr. J.J. Zhao for excellent technical and biosafety support. This work was supported by the National Key Research and Development Program (2020YFC0860100, 2020YFC0841401), the National Key Plan for Scientific Research and Development of China No.2016YFD0500306, the National Natural Science Foundation of China, China (82041006).

---

**Fig.S1.**
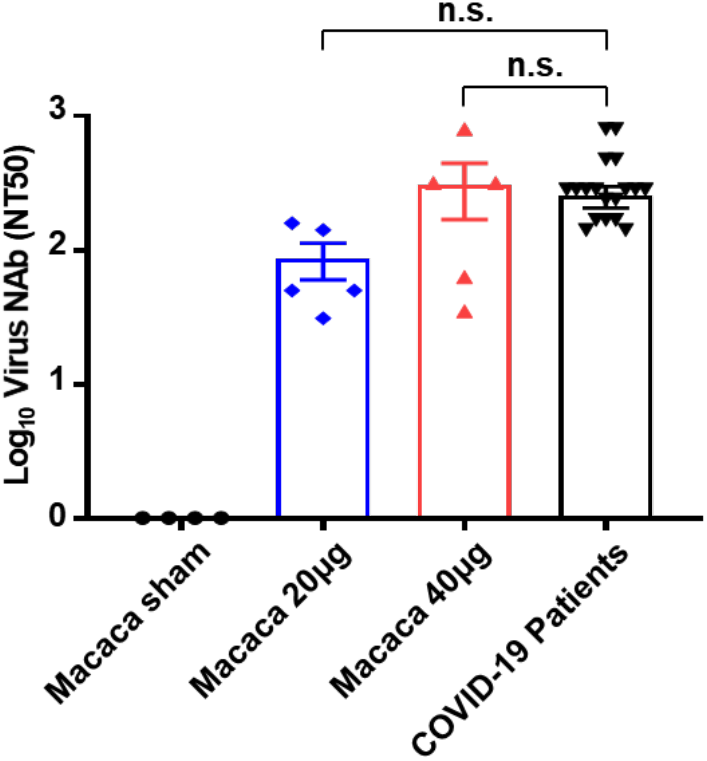
Comparison of neutralizing antibody titers in RBD-Fc-immunized monkeys and convalescent sera from COVID-19 patients. The serum neutralizing antibody titers were calculated from macaca fascicularis (n=4) immunized with 40 μg RBD-Fc and COVID-19 patients’ convalescent sera (n=23), respectively. Significance was calculated using a one-way ANOVA with multiple comparison tests. (n.s., not significant).

**Fig.S2.**
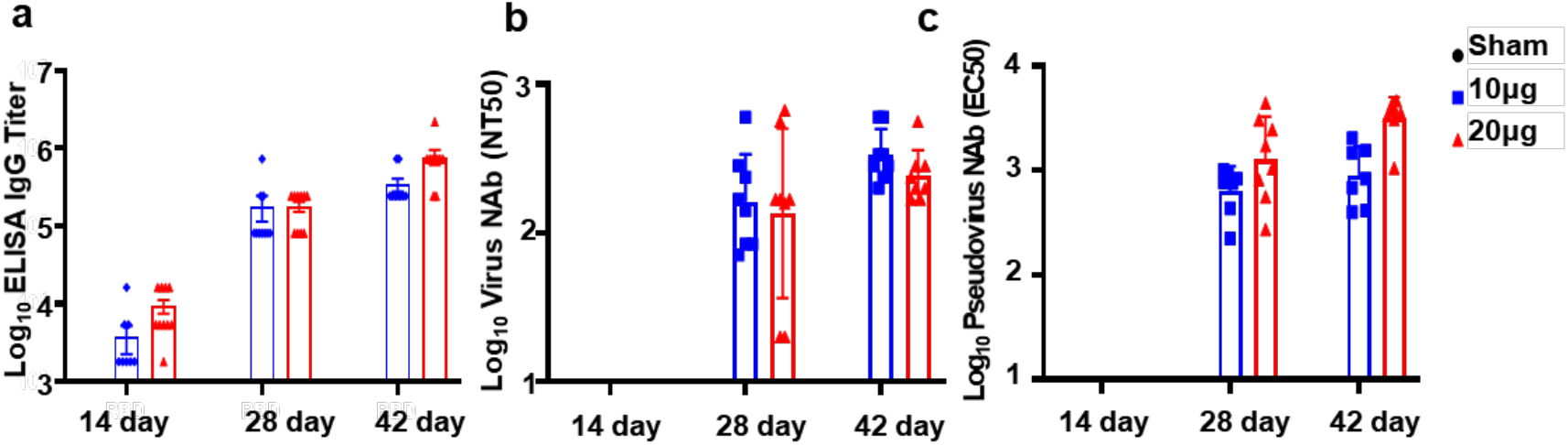
SARS-Cov2 RBD-Fc Vacc elicit strong humoral immunity in mice. hACE2-Tg mice were immunized on day 0, day 14, d28 with 10ug and 20ug doses of RBD-Fc Vacc or adjuvant and the serum were collected at the indicated time. a) The SARS-CoV-2 RBD specific IgG were examined by ELISA. b&c) Neutralizing antibodies were determined by microneutralization assay using both the SARS-CoV-2 (b) and pseudovirus (c).

**Fig.S3.**
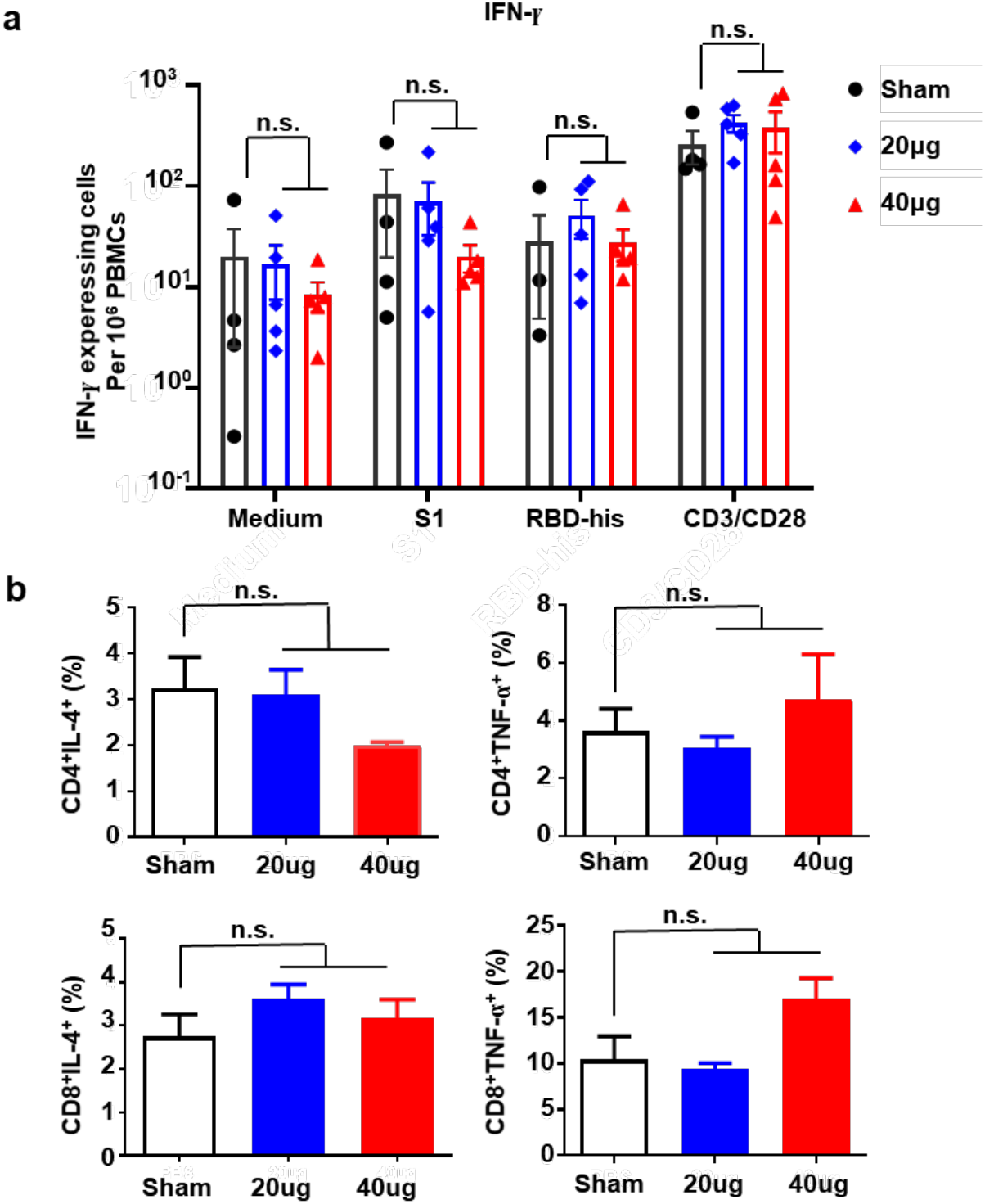
Characterization of the cellular immune response for SARS-CoV-2 RBD-Fc vaccine. **a)** ELISPOT assay for IFN-γ in PBMCs from macaca fascicularis. Data are shown as mean ± SEM. Significance was calculated using unpaired t-test (n.s., not significant). b). SARS-CoV-2 RBD-specific IL-4^+^T cells and TNF-a^+^T cells in CD4^+^T and CD8^+^T cells from macaca fascicularis PBMCs were detected by flow cytometry.

**Table S1.** The gender, the marked number for each animal and the Log10 gRNA copies/g values for each lung lobes of vaccinated or non-vaccinated macaca fascicularis were shown.

|  | Monkey<br>NO. | Gender | Left lobe |  |  | Right lobe |  |  |
| --- | --- | --- | --- | --- | --- | --- | --- | --- |
|  |  |  | Upper | Middle | Lower | Upper | Middle | Lower |
| Sham | 18123 | F | 7.68 | 8.47 | 5.87 | 7.48 | 6.04 | 8.24 |
|  | 17530 | M | 6.73 | 5.58 | 6.19 | 8.04 | 6.40 | 6.88 |
|  | 17497 | F | 6.69 | 8.92 | 9.67 | 4.79 | 4.57 | 4.72 |
|  | 18098 | M | 3.98 | 9.4 | 4.55 | 5.29 | 4.76 | 4.53 |
|  | average |  | 6.27 | 8.09 | 6.57 | 6.40 | 5.44 | 6.09 |
| 40ug | 17462 | M | 0 | 0 | 0 | 6.78 | 0 | 5.01 |
|  | 16022 | M | 0 | 4.38 | 0 | 3.56 | 3.95 | 3.94 |
|  | 17011 | F | 0 | 0 | 4.54 | 0 | 4.46 | 4.69 |
|  | 18071 | F | 0 | 4.41 | 0 | 0 | 0 | 5.41 |
|  | average |  | 0 | 2.20 | 1.14 | 2.59 | 2.10 | 4.76 |

**Fig.S4.**
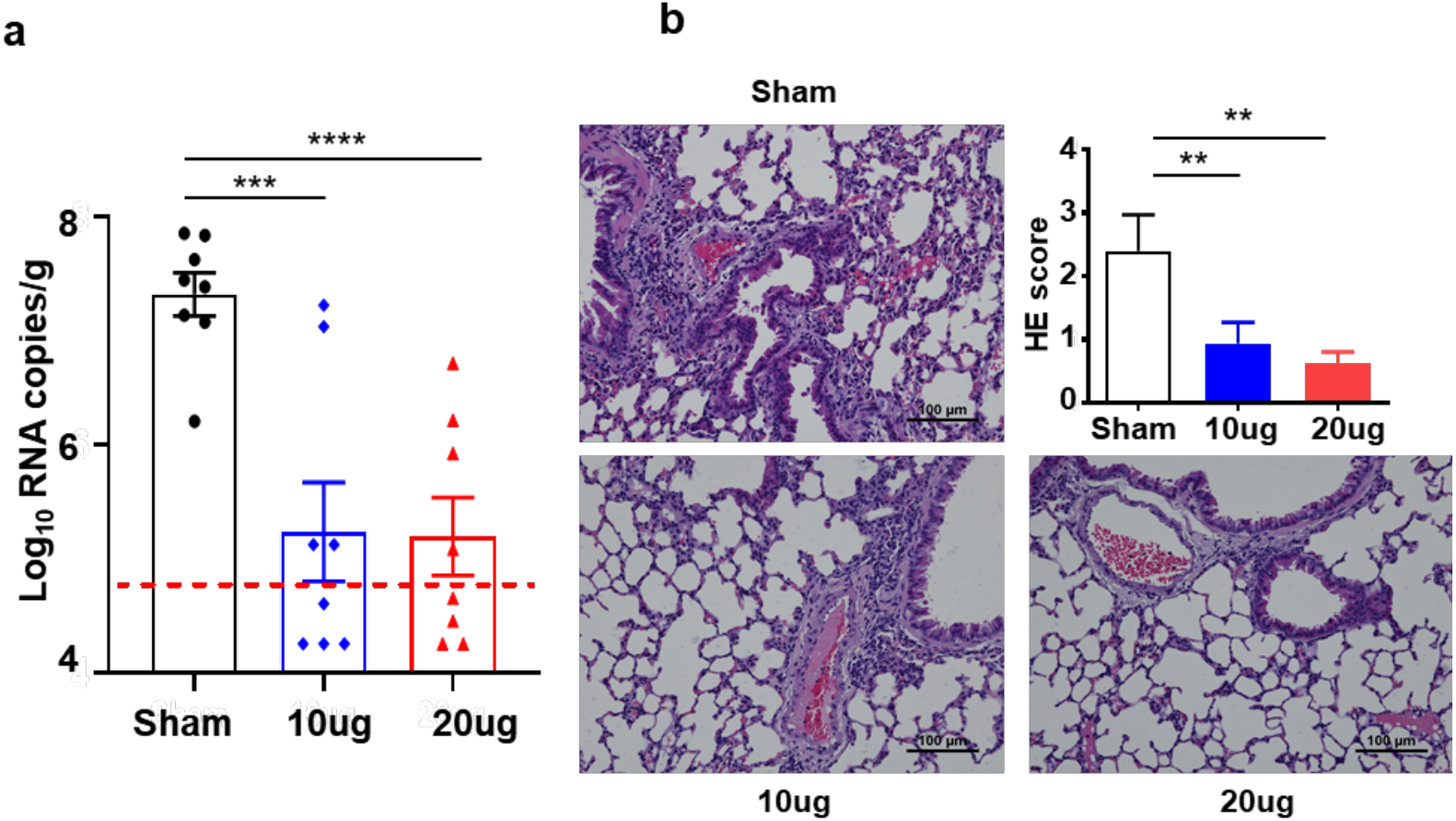
Protective efficacy of RBD-Fc Vacc in hACE2-Tg mice. hACE2-Tg Mice were immunized three times as described in the materials and methods (n=8). a) Viral loads in lung tissue from the inoculated mice at day 5 post infection were measured. All data presented as mean +SEM from four independent experiments. The dashed lines indicate the detection limit of the assay. b) Histopathological examinations in lungs from all the inoculated mice at day 5 post injection. Semi-quantitative analysis of histopathological changes of lung tissues. Data are presented as means ± SEM (n=6).

